# Lipid transport is necessary for neocortical lamination

**DOI:** 10.64898/2025.12.05.692589

**Authors:** Jacqueline Chrast, Stephan C. Collins, Catherine Roger, Siwar Ben-Ayache, Chiara Auwerx, Pauline Rogg, Fabrizio Vacca, Océane Musset, Lucie Gueneau, Yann Emmenegger, Fabienne Allias, Frederic Tran-Mau-Them, Daniel C. Koboldt, Matthew T. Pastore, Kristen V. Truxal, Linda Z. Rossetti, Lindsey Walker, Kiarash Sadrieh, Matthew A. Deardorff, Catarina Olimpio, Jessica Rosenblum, Marije Meuwissen, Anna C. Jansen, Jane Hassell, Manju Kurian, Ana Perez Caballero, Ben Paternoster, Oliver Klaas, Andreas Busche, Judit Horvath, Jean-Hubert Caberg, Serpil Alkan, Marijn F. Stokman, Stephanie Baskin, Nicolas Goss, Hector Gallart-Ayala, Muhammad Ansar, Zoltan Kutalik, Julijana Ivanisevic, Nicolas Guex, Binnaz Yalcin, Alexandre Reymond

## Abstract

We previously described the Alkuraya-Kučinskas syndrome, a disorder associated with biallelic variants in *BLTP1* (bridge-like lipid transfer protein), a.k.a. *KIAA1109*. The majority of probands die perinatally with corpus callosum agenesis, ventriculomegaly and arthrogryposis. Homozygous ablation of mouse *Bltp1* resulted in similar preweaning lethality. Here, we describe ten novel patients expanding the characterization of this syndrome at the mild end of the phenotypic spectrum. To model this syndrome, we engineered Emx1-Cre-mediated conditional knockouts (cKO) in which *Bltp1* expression is only removed in cortical and hippocampal neurons. This restricted ablation of *Bltp1* recapitulated the preweaning lethality observed in the constitutional knockouts, suggesting that lack of *BLTP1* expression in neurons is sufficient to cause death. Homozygous cKO presented a complete agenesis of the corpus callosum, a smaller anterior commissure, a malformed hippocampus and a reduced thickness of the cortical plate with a complete lack of defined structural layers and absence of radial glial and intermediate neural progenitors and mature neurons.

As BLTP1 was shown to be a barrel-shaped tube containing lipids, we compared the amount of lipid species in the cKO and their control littermates’ cortexes. We observed significant depletions of ether-linked phosphatidylethanolamines and triglycerides and accumulations of sphingomyelins and hexosylceramides in cKOs. Our results are consistent with the recent description of BLTP1 as a tubular protein that transports phospholipids between the endoplasmic reticulum and the plasma membrane. They suggest that non-vesicular lipid transport is essential for neocortical and cerebellar lamination.

Consistent with a BLTP1 role in cortex development we show that heterozygous carriers of a *BLTP1* truncation variant presented a decrease in peripheral cortical grey matter suggesting an autosomal dominant inheritance pattern beside the already described autosomal recessive.

## INTRODUCTION

Congenital anomalies are a major health issue affecting about 4% of newborns worldwide, leading to perinatal death(Gueneau et al. 2018; Oddsson et al. 2023; Mattioli et al. 2025) and life-long disabilities(Verma 2021). Whereas about half of the cases remain unsolved(Wojcik et al. 2023), a large fraction of these conditions has a monogenic genetic origin that could be identified by high-throughput sequencing techniques(Bamshad et al. 2019). Autosomal dominant, often *de novo*, variants represent the most common etiology in Western countries, whereas autosomal recessive inheritance is prevalent in countries with frequent parental consanguinity(Antonarakis 2021). We previously described the Alkuraya-Kučinskas syndrome (ALKKUCS; OMIM # 617822), a disorder that combines cerebral parenchymal underdevelopment and mild to severe ventriculomegaly with seizures, clubfoot and arthrogryposis, often accompanied by ophthalmic symptoms. It is caused by biallelic variants in *BLTP1* (bridge-like lipid transfer protein), a.k.a. *KIAA1109*(Gueneau et al. 2018), a gene chiefly expressed in the developing brain and cerebellum of vertebrates(Cardoso-Moreira et al. 2019). A large fraction of the 45 Alkuraya-Kučinskas syndrome individuals previously described died perinatally or were elected for premature termination of pregnancies (**Table S1**)(Gueneau et al. 2018; Kvarnung et al. 2018; Shamseldin et al. 2018; Filatova et al. 2019; Kane et al. 2019; Cabet et al. 2020; Kumar et al. 2020; Meszarosova et al. 2020; Chin et al. 2022; Yue et al. 2022; Gul et al. 2023; Rice et al. 2024; Turgut et al. 2024; Zhen and Li 2025). Homozygous ablation of mouse *Bltp1* similarly resulted in perinatal lethality, as well as abnormal embryo size(Skarnes et al. 2011; Liu et al. 2025).

Because of its structural similarity with VPS13A-D (vacuolar protein sorting 13 homolog A-D), the 5093 residue-long BLTP1 protein (NM_001384125.1) was suggested to form a hydrophobic tunnel that bridges the ER (endoplasmic reticulum) to the plasma membrane, enabling lipid transport(Jumper et al. 2021; Levine 2022; Hanna et al. 2023). Consistent with this hypothesis, (i) its yeast ortholog Csf1p was isolated in a screen for proteins important for inter-organelle lipid transport(John Peter et al. 2022) and was found at membrane contact sites, interacting with the ethanolamine phosphotransferase Mcd4p that adds the initial phosphoethanolamine to nascent GPI in the ER(Toulmay et al. 2022); (ii) its *D. melanogaster* ortholog TWEEK was shown to be necessary for synaptic vesicle recycling and regulation of neuromuscular junction (NMJ) size(Verstreken et al. 2009; Khuong et al. 2010); (iii) the knockdown of its *C. elegans* orthologous gene, *lpd-3*, led to a lipid-depleted phenotype and abnormal phospholipid distribution(McKay et al. 2003; Wang et al. 2022); (iv) knockdown of *BLTP1* in HEK293 cells and ablation of *Bltp1* in mouse fibroblasts perturbed the distribution of PIP3 phospholipids(Wang et al. 2022); (v) mouse embryos homozygously ablated for *Bltp1* presented with smaller NMJ(Liu et al. 2025); and (vi) cells of Alkuraya-Kučinskas syndrome patients presented with defects in endosomal trafficking(Kane et al. 2019). Further confirmation came from the experimentally solved structure of the first ∼1,700 residues of LPD-3, which showed a rod-shaped tunnel filled with ordered phospholipid molecules(Kang et al. 2025). LPD-3 co-purified with Spigot, the human ortholog of C1orf43, and Tmem-170, the ortholog of the TMEM170A/B scramblases(Kang et al. 2025; Rocha-Roa et al. 2025).

Here we expand the phenotypic spectrum of Alkuraya-Kučinskas syndrome by describing ten additional patients and show that homozygous ablation of *Bltp1* in mouse cortical and hippocampal neurons recapitulates the preweaning lethality of the *Bltp1* constitutive knockout. These brain-conditional knockouts presented with severe disruption of neocortical lamination and significant alterations of the cortex lipidome, i.e. an increase in sphingolipids and a depletion of ether-linked phosphatidylethanolamines and triglycerides

## RESULTS

### Alkuraya-Kučinskas syndrome

Published Alkuraya-Kučinskas syndrome individuals presented with malformations of the central nervous system, as well as associated craniofacial abnormalities affecting the ears, eyes, and mouth (**Table S1**)(Gueneau et al. 2018; Kvarnung et al. 2018; Shamseldin et al. 2018; Filatova et al. 2019; Kane et al. 2019; Cabet et al. 2020; Kumar et al. 2020; Meszarosova et al. 2020; Chin et al. 2022; Yue et al. 2022; Gul et al. 2023; Rice et al. 2024; Turgut et al. 2024; Zhen and Li 2025). Using data aggregation of multiple laboratories and clinical centers, e.g. GeneMatcher(Sobreira et al. 2015), we identified ten new patients from nine families with biallelic variants in *BLTP1* (**Table S2**).

A large fraction of the 45 Alkuraya-Kučinskas syndrome individuals previously described(Gueneau et al. 2018; Kvarnung et al. 2018; Shamseldin et al. 2018; Filatova et al. 2019; Kane et al. 2019; Cabet et al. 2020; Kumar et al. 2020; Meszarosova et al. 2020; Chin et al. 2022; Yue et al. 2022; Gul et al. 2023; Rice et al. 2024; Turgut et al. 2024; Zhen and Li 2025) and of the ten patients identified in the present report, presented with perinatal death or were elected for premature termination of pregnancies (**Tables S1-S2**) (67%, 37 out of 55 cases). Specifically, all affected individuals carrying biallelic truncating variants died perinatally (21/23; 91%) with the exception of a mother and her daughter, who carry a homozygous noncanonical splice donor variant suggested to lead to less severe consequences via leaky splicing (**Table S1**)(Chin et al. 2022). Affected individuals compound heterozygote for a truncation and a missense variant and affected individuals with biallelic missense variants in *BLTP1* exhibited lower rates of perinatal death (5/11 (45%) and 11/21 (52%), respectively). To model the possible effect of the missense variants identified in published and novel Alkuraya-Kučinskas syndrome cases (**Tables S1-S2**), we used the experimentally solved cryogenic electron microscopy (cryo-EM) structure of the first ∼1,700 residues of LPD-3(Kang et al. 2025), in addition to the human AlphaFold model(Jumper et al. 2021). Whereas their overall channel fold is similar, BLTP1 has longer loops than LPD-3 (**Figure 1**). Their structure consists of 17 repeats of β-groove (RBG), typically composed of five anti-parallel strands and one helix connected by a long loop(Levine 2022). While the BLTP1 longest isoform is 5,093 residues long, about two-thirds of the missense variants reported here and previously(Gueneau et al. 2018; Kvarnung et al. 2018; Shamseldin et al. 2018; Filatova et al. 2019; Kane et al. 2019; Cabet et al. 2020; Kumar et al. 2020; Meszarosova et al. 2020; Chin et al. 2022; Yue et al. 2022; Gul et al. 2023; Rice et al. 2024; Turgut et al. 2024; Zhen and Li 2025) (15 out of 23, 65%) are located within its first 1,700 residues (**Tables S1-S3**). The predicted effects of the missense variants are detailed in **Table S3,** and some examples are shown in **Figure 1A**. Eight of the 23 missense variants were identified in individuals who died perinatally suggesting that the activity of the encoded protein is severely impaired by analogy with the perinatal death associated with truncation variants. Seven of these variants, N216K, D228G, P433A, T2828P, P3050H, G3385R and A4333T (A4245T in Q2LD37), are positioned within beta-strands that form the RBG repeats barrel; the exception is R968C, which is in a well-structured region flanking the barrel (**Figure 1**). The variants K957T and P1631R are likewise positioned within the barrel. Some of the identified variants affect residues in regions interacting with Spigot/C1orf43 (Y47C, N216K and D228G) and/or in contact or neighboring residues in contact with lipids (Y47C, G437E and L855P) (**Figure 1**). The remaining variants are in unstructured loops with (D761N, T811A, P979S and V1867M) or without a *C. elegans* counterpart (Y1329S, M1573I, R1958Q, H2246R, C4317S). F1556S is in an anti-parallel beta-strand that folds perpendicularly to the exterior of the main RBG barrel, a fold conserved in *C. elegans*. The G3385R variant was identified in three Tunisian families suggesting a possible founder effect (families F5, F6 and F14) (**Table S1**)(Gueneau et al. 2018; Cabet et al. 2020).

**Figure 1.**
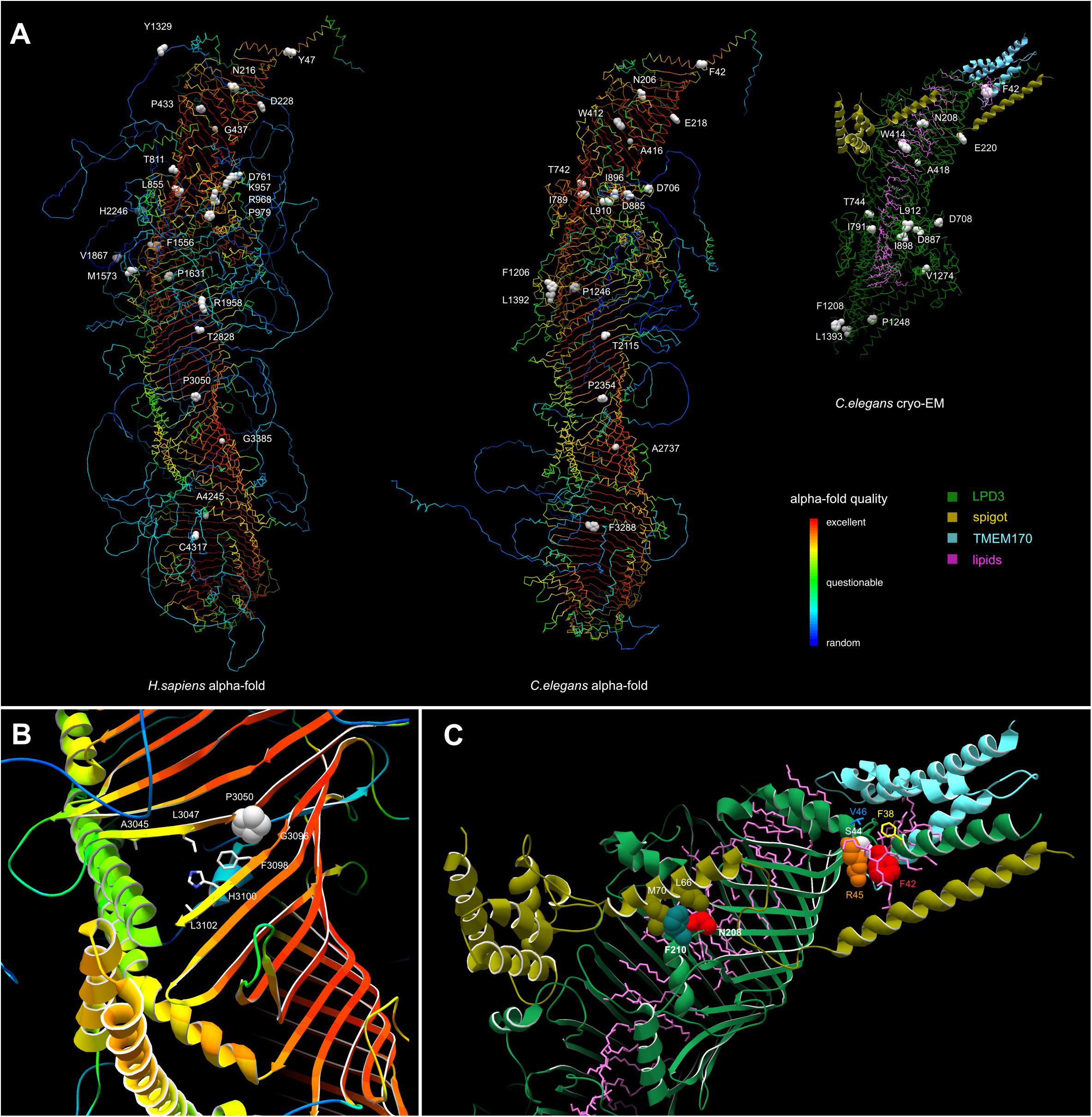
BLTP1 missense variants. (**A**) **Left:** AlphaFold model of human BLTP1, colored by increasing percent confidence of the model (light blue, dark blue, green, yellow, orange, red). Position of the missense variants identified in Alkuraya-Kučinskas syndrome individuals are indicated. **Center:** AlphaFold model of the *C. elegans* ortholog (LPD-3), taken from reference(Pandey et al. 2023) highlighting the position of residues corresponding to the human variants. Same color code as in left vignette. **Right:** Cryo-EM structure of the *C. elegans* ortholog (LPD-3) (Kang et al. 2025) highlighting the position of residues corresponding to the human variants. Dark green ribbon: LPD-3; asparagus ribbon: Spigot; cyan: TMEM170; purple: lipids; **(B)** P3050 is positioned within the RBG repeats barrel. It changes the orientation of a strand, which disrupts the continuous structure of the beta-sheet and leaves space for a passage in/out of the beta-barrel between the strands P3043-L3049 and the strand F3098-L3102. Both the residue and its associated kink are structurally conserved in the *C. elegans* AlphaFold model (P2354). Same color code as in left and central vignettes of panel A. (**C)** *C. elegans* cryo-EM PDB 9CAP. Dark green ribbon: LPD-3; asparagus ribbon: Spigot; purple: lipids. **Left:** Context of the N216K variant using the corresponding N208 residue of *C. elegans*, which is in a region critical for the proper interaction with a Spigot H1 alpha-helix. Spigot conserved L66 (asparagus spacefill) residue contacts N208 (red spacefill), whereas F210 (equivalent to Y218 in human; teal spacefill) interacts with Spigot M70 sidechain (Q in human ortholog C1orf43; asparagus spacefill). **Right:** Context of the Y47 variant using the corresponding F42 residue (red spacefill) of *C. elegans*; S44 (structurally equivalent to human S49): white spacefill; R45 (structurally equivalent to human R50): orange spacefill; V46 (structurally equivalent to human N51): blue stick; F38 (structurally equivalent to human Y43): yellow stick.

Brain anomalies described in Alkuraya-Kučinskas syndrome individuals included hydrocephalus, cerebellar anomalies (cerebellar hypoplasia and dysplasia of the cerebellum) and dysgenesis of the corpus callosum (corpus callosum agenesis, a shortened corpus callosum and hypogenesis of the corpus callosum) in 76% (34 out of 45), 63% (26/41) and 59% (26/44) of cases, respectively. Additionally, 47% (18/38) of affected individuals exhibited parenchymal anomalies including cerebral parenchymal underdevelopment and thinning. Lissencephaly was observed in 33% (12/36) of the cohort, and anomalies of the brainstem were present in 31% (11/36) including brainstem dysgenesis, dysplasia of the brainstem, kinked brainstem and Z-shaped appearance of the brainstem. Pons alterations were observed in 21% (7/34) of the affected individuals (hypoplasia, flattening, elongation, and abnormal segmentation). Cortical malformations were seen in 18% (6/34) of cases, including cortical dysplasia, cortical thinning, and absence of cortical lamination (**Figure 2A**). Cerebellar vermis phenotypes, including atrophy, dysgenesis, and reduction in size, were identified in 18% (6/34) of cases. Lastly, other white matter malformations (excluding anomalies related to the corpus callosum) were present in 9% (3/34) of cases.

**Figure 2.**
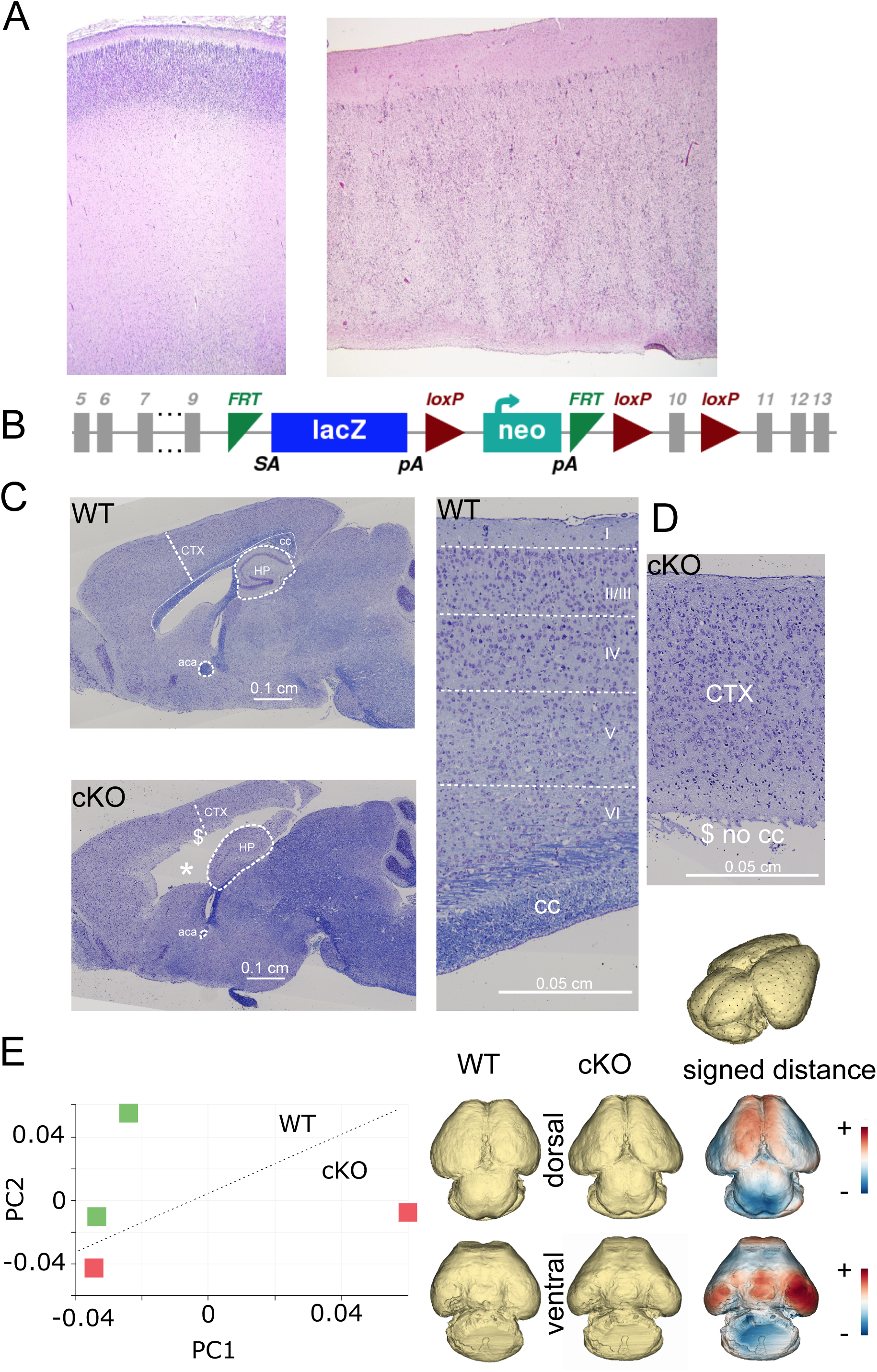
Neuroanatomical studies implicate *Bltp1* in cortical development and brain shape. (**A**) Cortical section of 21 weeks control (**left**) and Alkuraya-Kučinskas syndrome patient (right) fetuses. Note the unlaminated cortex of the Alkuraya-Kučinskas syndrome patient. (**B**) IMPC-generated allelic construction which contains a LacZ trapping cassette and a floxed promoter-driven neomycin cassette inserted into intron 9 of *Bltp1* with loxP sites flanking exon 10. (**C**) Representative Nissl-Luxol double-stained parasagittal brain sections at Lateral +0.72 mm from adult male *Bltp1* heterozygous (**left**) and homozygous (**right**) conditional knockout mice, with *Bltp1* selectively inactivated in the Emx1-expressing lineage. The heterozygous mouse serves as a control, due to the recessive inheritance pattern of the condition and the unavailability of littermate WT mice. Scale bar represents 0.1 cm. The dollar sign ($) indicates agenesis of the corpus callosum, and the asterisk (*) indicates enlarged lateral ventricles. (**D**) Magnified views highlighting the cortical phenotype, with no visible layering in the homozygous cKO (Emx1^Cre^*; Blpt1*^fl/fl^; **right**) compared to the heterozygous control (**left**), where cortical layers (I to VI) are visually distinguishable. Scale bars are 0.05 cm for both the right and left images. Annotations: CTX (cortex), HP (hippocampus), aca (anterior commissure), cc (corpus callosum). (**E**) Morphogeometric analysis of E18.5 cKO mouse brains using pseudolandmarks and PCA-based deformation modeling. Three-dimensional models were analyzed using 3D Slicer and SlicerMorph. A combination of fixed anatomical landmarks and pseudolandmarks (generated with the PseudoLMGenerator) was used to capture overall brain surface morphology. Landmark configurations were aligned using Generalized Procrustes Analysis (GPA), and principal component analysis (PCA) was performed on the resulting shape data. The models shown represent PCA extrapolations along the first two principal components, illustrating the major axes of morphological variation between WT and cKO groups. Model-to-model distance maps quantify regional differences between WT and cKO brains: red regions indicate areas where distance increases (expansion in cKO relative to WT), and blue regions indicate areas where distance decreases (contraction in cKO relative to WT).

While the majority of the ten novel affected individuals with bi-allelic variants in *BLTP1* present with various cardinal characteristics of the syndrome (patients no. 46-52), we also uncovered three patients with milder symptoms (**Table S2**) akin to previously described families(Kumar et al. 2020; Chin et al. 2022). Patients no. 53-55 showed autistic traits and developmental delay largely without syndromic features and no overt brain anomalies suggesting extensive variable expressivity of Alkuraya-Kučinskas syndrome (**Table S2**).

### Conditional Bltp1 mouse model

The IMPC (International Mouse Phenotyping Consortium) reported that homozygous ablation of *Bltp1* results in a fully penetrant pre-weaning lethality(Skarnes et al. 2011), which was confirmed in a recent study(Liu et al. 2025). Whereas reminiscent of the incompatibility with life observed in humans carrying loss-of-function variants in the *BLTP1* ortholog, it does not help deconvolute tissue-specific effects brought about by ablation of this gene. To create conditional tissue-specific knockouts (cKO), we established a colony of *Bltp1*^tm1a/+^ mice, which carry a LacZ trapping cassette and a floxed promoter-driven neomycin cassette inserted into the intron 9 of *Bltp1* and loxP sites flanking its exon 10 (**Figure 2B**). We crossed them with transgenic mice ubiquitously expressing Flp recombinase to convert the tm1a allele into the conditional tm1c allele with loxP sites on either side of exon 10. *Bltp*1^tm1c^ mice were then crossed with (B6.129S2-Emx1^tm1(cre)Krj/J^) mice that express Cre recombinase under the control of the Emx1 (empty spiracles homeobox 1) promoter to specifically ablate *Bltp1* in neurons by conditionally removing the floxed exon and generating a tm1d allele. The detailed crossing scheme is presented in **Figure S1**.

The specific homozygous deletion of *Bltp1* exon 10 in cortical and hippocampal neurons significantly decreased survival rates compared to wildtype (WT) mice (one observed versus 21 expected: 27 tm1c^+/-^ mice (33%), 33 tm1c^-/-^ mice (40%), 22 tm1d^+/-^(Emx1^Cre^*; Blpt1*^+/fl^) mice (27%) and one tm1d^-/-^ (Emx1^Cre^*; Blpt1*^fl/fl^) mouse (1%), N_total_=83, p=3.65×10^-06^). Our results suggest that inactivation of *Bltp1* within the Emx1-expressing lineage is sufficient to recapitulate the mortality observed in constitutional knockouts of that gene.

### Bltp1 conditional knockout brain anomalies

We measured 22 brain structures totaling 40 measurements at Lateral +0.60 mm in the single Emx1^Cre^*; Blpt1*^fl/fl^ mouse (tm1d^-/-^ allele, referred as cKO hereafter) mouse that survived to adulthood using our described pipeline(Collins et al. 2018; Collins et al. 2019) (**Table S4**). Although the total brain area was similar between the cKO and WT mice, we identified previously unreported neuroanatomical phenotypes, including an 80% reduction in the anterior commissure and a malformed hippocampus (**Figure 2C**, bottom panel). More specifically, we noted disturbed lamination of the hippocampus, with no clear delineations between the Cornu Ammonis (CA) layers and the dentate gyrus, a pattern reminiscent of neuronal migration disorders(Belvindrah et al. 2014). Based on these observations, we hypothesize that individuals with Alkuraya-Kučinskas syndrome might also present with undetected hippocampal malformations. We also observed a complete agenesis of the corpus callosum (indicated by the dollar sign in **Figure 2C**), a severe enlargement of the lateral ventricles (indicated by an asterisk in **Figure 2C**), and a reduced cortical thickness with complete absence of cortical lamination (**Figure 2D**). These neuroanatomical phenotypes were similar to those observed in patients (**Figure 2A** and **Table S1**).

Next, we interrogated the cause of the observed cortical phenotype by studying brain morphology at embryonic day 18.5 (E18.5), using newly developed 3D histology approaches in combination with our previously published method for neuroanatomical quantification of embryonic brains based on 2D histology(Nguyen et al. 2022). Our 3D histological pipeline is based on high-resolution episcopic microscopy (HREM). In this study, morphogeometric analysis was used to assess brain shape at E18.5. Although the number of replicates was limited, we observed whole-brain shape differences along the dorsal axis in cKO embryos. The posterior part of the cortex appeared contracted, while the medial region was expanded, suggesting that the underlying neuroanatomical phenotypes may also influence overall brain shape (**Figure 2E**). Our 2D histological approach, based on the sagittal section at Lateral +0.60 mm, covered 17 developmentally distinct brain regions for 40 quantifiable brain parameters of area, height and width measurements (**Table S5-S6** and **Figure3A-C**). Prior to image analysis, each brain section underwent rigorous quality control, ensuring accurate stereotaxic positioning and symmetry along both dorsoventral and rostro-caudal axes. The neuroanatomical profile of both male and female cKO mice showed overall similarities, with the size of assessed structures appearing normal, made exception of the cortical plate. As hippocampal and corpus callosum development in mice occurs mainly postnatally, we speculate that the earliest phenotype manifests at the cortical level, followed by hippocampal lamination defects and corpus callosum agenesis, consistent with our observations in adult mice (**Figure 2C**). The cortical plate exhibited thinning (**Figure 3D-E**), consistent with the adult phenotype (**Figure 2D**). However, a pronounced difference was observed in the cortical plate thickness between sexes. Male embryos displayed a significant reduction of 39% (p=0.023), while the reduction in female embryos was comparatively milder at 17% (p=0.0068; **Figure 3E**).

**Figure 3.**
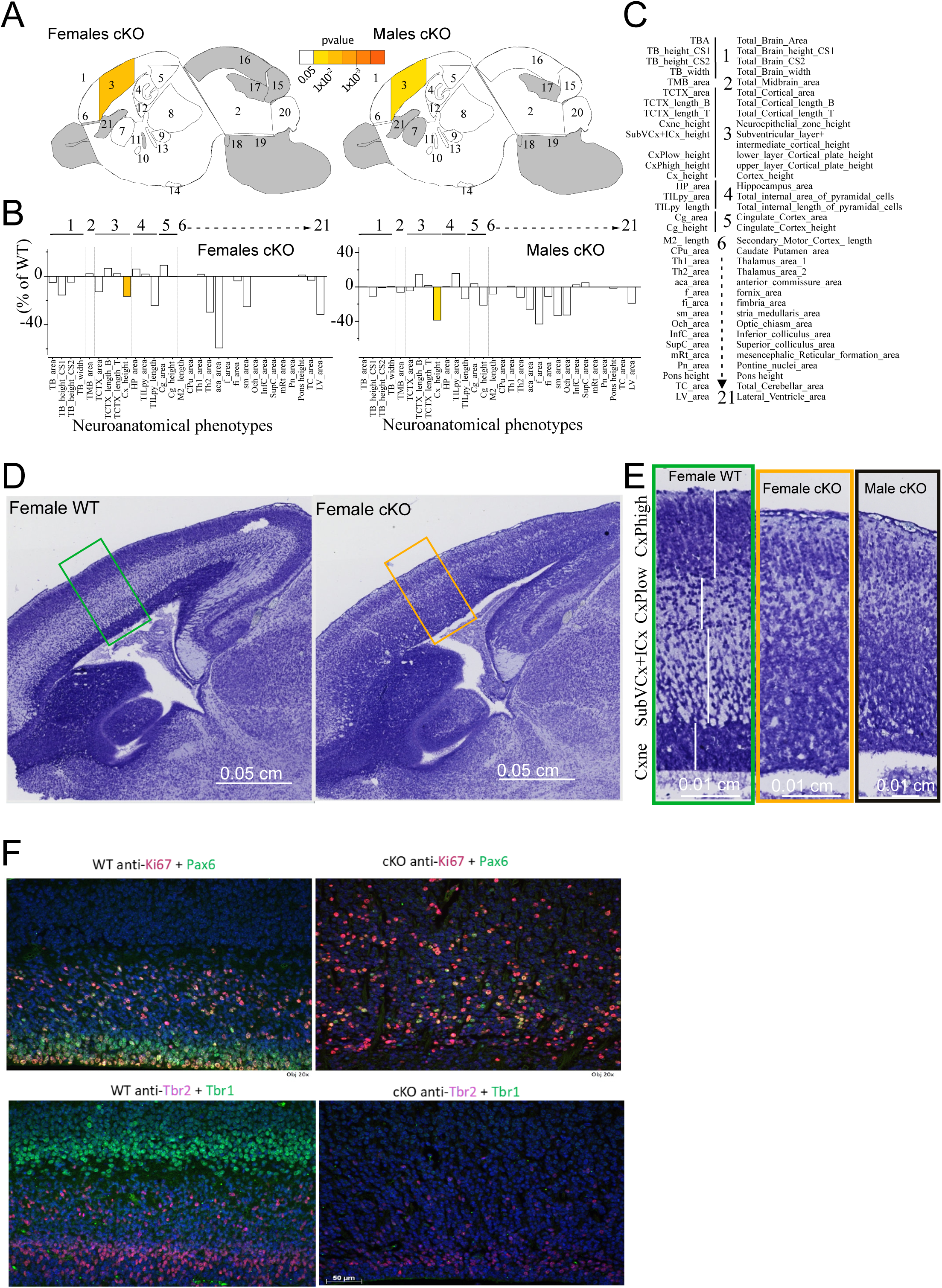
*Bltp1* plays a key function in the lamination process of the cerebral cortex. (**A**) Schematic illustration of a mouse sagittal brain section at embryonic day 18.5 (E18.5). Numbers identify the different brain regions assessed for area and length differences between cKO (*Emx1-Cre; Bltp1^fl/fl^*) and control littermates in females (**left**) and males (**right**). The light and dark yellow shading highlights the statistically significant differences (from p<0.05 to p<10^-4^, respectively) while the grey shading denotes regions not analyzed for this study. (**B**) Histograms depicting the size changes observed in female (**left**) and male (**right**) cKO embryos relative to littermate control embryos (set as 0) for each of the measured parameters (listed in **Tables S5-6** and on the bottom right-hand side of the Figure). (**C**) List of parameters measured with full name. Statistical analyses were performed with R, using two-tailed Student’s *t*-tests of equal variances. Due to cortical disorganization in conditional knockout mice, Cx_height corresponds to the sum of all cortical layers. (**D**) Representative images of female control (WT) and cKO sagittal brain sections (Lat +0.60 mm) stained with Luxol-Nissl. Insert is detailed in panel D. Scale bars: 0.05 cm. (**E**) Representative images of cortical layering in WT and structural layer disorganization in cKO mice. Scale bars: 0.01 cm. (**E**) List of parameters measured with full name. Statistical analyses were performed with R, using two-tailed Student’s *t*-tests of equal variances. Due to cortical disorganization in conditional knockout mice, Cx_height corresponds to the sum of all cortical layers. (**F**) Representative immunofluorescence images of KI67 (red) and Pax6 (green) (**top panels**) and Tbr2 (red) and Tbr1 (green) (**bottom panels**) of E18.5 WT (**left panels**) and cKO (**right panels**) sagittal cortical sections.

We could not determine the boundaries between the neuroepithelial (Cxne), subventricular (SubVCx), intermediate (ICx), lower layer of the cortical plate (CxP_low) and upper layer of the cortical plate (CxP_high) in cKO embryos. To help discern which layer(s) contributes to the thinning of the cortex (**Figure 3E**), we used markers for radial glial progenitors (Pax6), intermediate neural progenitors (EOMES/Tbr2), mature neurons (Tbr1), dividing (Ki67) and apoptotic cells (activated caspase 3) to identify the missing cells/layers(Kee et al. 2002; Englund et al. 2005). We observed almost no radial glial and neural progenitor cells and a total absence of post-mitotic neurons in cKO brains but detected many dividing cells throughout the thinner cKO cortex, made exception of the Cxne (**Figure 3F**). In contrast, whereas apoptotic cells were present in WT they were almost totally absent from cKO cortexes. A similar absence of lamination has been pinpointed in some Alkuraya-Kučinskas syndrome fetuses (**Figure 2A**), suggesting conserved neurodevelopmental principles between humans and mice. Together, our findings indicate that *Bltp1* plays a major role in the development and structural organization of the cerebral cortex.

### Heterozygous variants

GnomAD v4.1.0(Chen et al. 2024) population metrics indicate that *BLTP1* is under constraint for both truncation (loss-of-function) (pLOEUF=0.58) and missense variants (Z_missense_=7.69), suggesting that *BLTP1* heterozygous variants decrease fitness. Correspondingly, IMPC reports that constitutive heterozygous *Bltp1*^+/-^ mice present with abnormal behavior, decreased locomotor activity, increased thigmotaxis and startle reflex and abnormal lens/retina morphology(Skarnes et al. 2011). We thus examined whether hemizygosity of *Bltp1* in neurons could lead to disruptions in cortical lamination using the histological methods described above. Whereas we found no overt neuroanatomical phenotypes in heterozygote cKO embryos (**Table S5-S6, Figure S2**), we found that UK Biobank participants heterozygous for a *BLTP1* truncation variant presented a Bonferroni-significant decrease in peripheral cortical grey matter normalized for head size (N=183 carriers; N=166 controls; p=0.00872). An autosomal dominant inheritance of *BLTP1* variants is further substantiated by the published identification of eleven carriers of *de novo* missense variants and one carrier of a *de novo* stop gain variant but no carriers of *de novo* synonymous variants in *CSMD2* in 31,058 parent-offspring trios of individuals with developmental disorders (Kaplanis et al. 2020).

### Bltp1^-/-^ Lipids

As LPD-3 was shown to be a barrel shape tube containing lipids(Kang et al. 2025) and as BLTP1 and its orthologs were suggested to act as a bridge that mediate the traffic of lipids from the ER to both the plasma membrane(Wang et al. 2022) and mitochondria(Toulmay et al. 2022), we quantified lipid molecular species in the cortex of cKO embryos (four females and five males) and compared them to the amounts found in their control littermates (five females and seven males, N_total_=21; **Figure S1**). We quantified 394 lipids and identified 66 and 108 species significantly changed between cKO and controls at adjusted p-value cutoffs of <0.01 and <0.05, respectively (**Table S7** and **Figure 4A**). Hierarchical clustering shows an increase in the level of several sphingolipid species, including sphingomyelins (SM), ceramides (Cer) and hexosylceramides (HexCer), as well as a depletion of multiple triglycerides (TG) and specific species of phosphatidylethanolamines (PEs), especially ether-linked PEs (e-PE) in cKOs. Several depleted PEs were polyunsaturated and many contained polyunsaturated fatty acids (PUFA), such as C22:6 (docosahexaenoic acid (DHA)), C22:5 (docosapentaenoic acid (DPA)) and C20:4 (arachidonic acid (AA)), that are known to play an important role in maintaining the membrane fluidity, elasticity and flexibility (**Table S7**).

**Figure 4.**
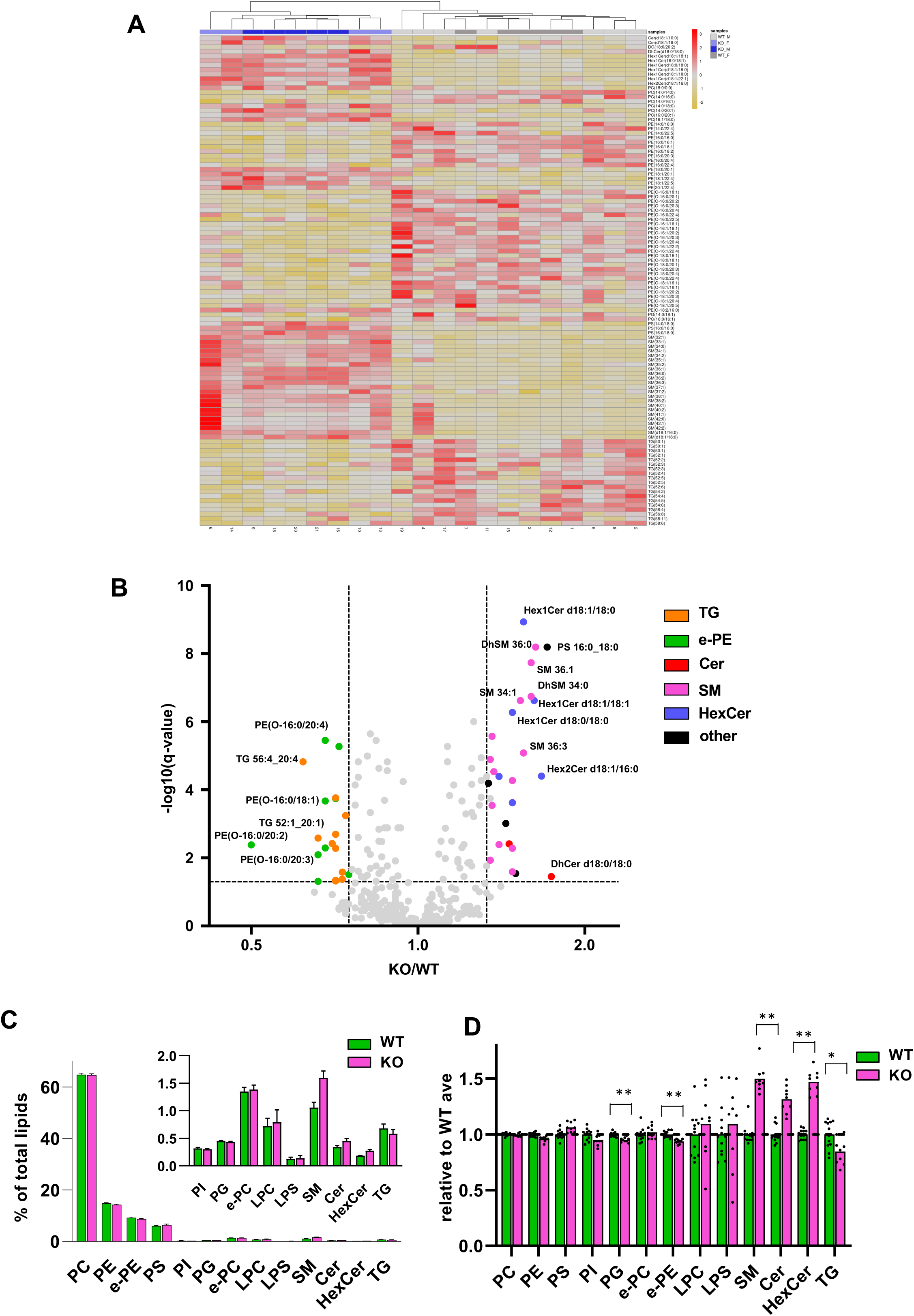
Lipid profile of E18.5 cortexes ablated for *Bltp1*. (**A**) Hierarchical clustering heat map representation of the levels of the 108 lipids species significantly changed at an adjusted p-value cutoff of <0.01 in cortexes of E18.5 cKO (*Emx1-Cre; Bltp1^fl/fl^*) (columns 1-9: four females light blue and five males dark blue) compared to E18.5 WT controls (columns 10-21: five females light grey and seven males dark grey) ranked by lipid class. In cKO, we observed a depletion of specific species of ether-linked phosphatidylethanolamines (e-PE) and triglycerides (TG) and an accumulation of sphingomyelins (SM), ceramides (Cer) and hexosylceramides (Hex1Cer) species (**B**) Volcano plot illustrating relatives changes in individual lipid species in cKO (N=9) relative to WT (N=12). **(C)** Absolute molar abundance of lipid classes expressed as percentage of total lipid content in cKO cortexes. (**D**) relative changes in lipid classes relative to the corresponding WT average level. (* q<0.05; **q<0.01).

As sample clustering indicates that sex as a negligible effect on the lipid composition compared to genotype, we only consider the latter in our subsequent analyses, e.g. lipidome volcano plot comparing WT and cKOs (**Figure 4B**). We then compared overall lipid classes alteration by summing molar amounts of the individual lipid species (**Figure 4C**). We observe a significant increase in the amount of all three sphingolipid classes (q<0.01) ranging from a 51% increase in SM to a 32% increase in Cer (**Figure 4D**). It stems from an increase of almost all sphingolipid species leaving unaltered species distribution within each class rather than from differences among individual species (**Figure S3**). Compared to the increase in sphingolipid, the class analysis evidences a significant (q<0.01) albeit less important decrease in total e-PE content (6% decrease), which results from the decrease of specific e-PE species (**Figure 4D and S4**). Upon inspection all decreased species present alkyl chains with one or no insaturation in the sn-1 position while species with two insaturations in sn-1 are generally increased or unaltered. We also observe significant decreases in total TG (-15%, q<0.05) and PG contents (-5%, q<0.01) (**Figure 4D**). Our Lipid ontology analyses(Molenaar et al. 2019) identified enrichment of “lipid storage” and “lipid droplets”, two terms related to sphingolipid accumulation and TG depletion respectively, of “mitochondrion” and “negative intrinsic curvature”, two terms associated with PE depletion as well as “plasma membrane” and “high transition temperature”, which are linked to SM accumulation (**Figure 4B**).

## DISCUSSION

We show that mice homozygously ablated for *Bltp1* only in cortical and hippocampal neurons present with almost completely penetrant preweaning lethality akin to what is observed in the constitutional knockout of that gene. These results suggest that, contrary to what was previously suggested, the cause of death of *Bltp1^-/-^* mice might not be respiratory failure(Liu and Lin 2022) but instead due to disruption of lipid transport in the brain. Similar to some Alkuraya-Kučinskas syndrome individuals, Emx1^Cre^*; Blpt1*^fl/fl^ mice presented complete agenesis of the corpus callosum, malformed hippocampus, decreased anterior commissure and reduced thickness of the cortical plate with complete absence of lamination. Our results also suggest that the statement that lamination is not necessary for cognition should possibly be revisited (Guy and Staiger 2017; Lossi et al. 2019).

Common variants in the *BLTP1* locus were associated with blood measurements (mean platelet volume, mean reticulocyte volume, serum total protein levels) and type 1 diabetes by genome-wide association studies (GWAS)(Sollis et al. 2023). Correspondingly, IMPC constitutive *Bltp1^+/-^* mice presented with impaired glucose tolerance and increased circulating glucose level(Skarnes et al. 2011). The *BLTP1* locus was also linked to immune reaction and autoimmunity phenotypes, as well as mood disorders. There was also an impact on quality of life as the *BLTP1* locus was also associated with diminished household income and educational attainment(Sollis et al. 2023). These associated traits might explain the significant paucity of truncation and missense variants indicated by gnomAD population metrics. Here, we correspondingly expand the phenotypic features associated with rare *BLTP1* variants beside enrichment in parent-offspring trios of individuals with developmental disorders (Kaplanis et al. 2020). Firstly, heterozygous carriers of truncation variants present a cortical phenotype. Secondly, while the vast majority of carriers of bi-allelic *BLTP1* variants present with brain anomalies and die perinatally (**Table S1-S2**), we identified three affected individuals with mild disease presentations consisting primarily of non-syndromic developmental delay. Importantly, we cannot rule out that these predicted-to-be deleterious variants are spuriously associated and not causative of the observed phenotypes. Future studies and the identification of more patients are needed to understand if the hypothesis of causativeness of these *BLTP1* variants is correct.

Our lipidome profiling shows severe lipid alterations in the cortex of Emx1^Cre^*; Blpt1*^fl/fl^ mice. Our findings align with the recently published cryo-EM structure of LPD-3 showing a tube-forming lipid transport protein (**Figure 1**)(Kang et al. 2025) and the suggested lipid transfer role of BLTP1 (Jumper et al. 2021; Levine 2022; Hanna et al. 2023). Its ablation may result in defects in intracellular lipid transport processes that may lead to altered membrane bilayers composition impairing its biophysical and signaling properties impacting neuronal differentiation and lamination in the cortex. Correspondingly, the BLTP1 yeast and worm orthologs were shown to be important in the regulation of biophysical membrane properties such as homeoviscous adaptation and resistance to cold(John Peter et al. 2022; Wang et al. 2022). Many of the PEs depleted in Emx1^Cre^*; Blpt1*^fl/fl^ mice are ether-linked PEs, which are suggested to facilitate membrane fusion processes, as well as to stabilize lipid rafts(Sublette et al. 2024). Some of the BLTP1-associated traits, i.e. bipolar disorder, schizophrenia, major depressive disorder and suicide risk, have been associated with a decrease of PUFA levels(Baccouch et al. 2023). Further supporting the role of PUFA in these pathologies, we found that these fatty acids are components of the PE species depleted in the cortex of Emx1^Cre^*; Blpt1*^fl/fl^ mice.

Is the lack of BLTP1 and the resulting diminished transport of some classes of lipids at the root of the observed lipidome differences? We observe an important increase in sphingolipids with a more selective depletion in PE, e-PE and TG. Given the absence of lamination and the associated drop in cell diversity it is intrinsically difficult to establish which lipid alterations could primarily be dependent of BLTP1 loss, and which are secondary to the associated developmental defects. The initial steps of ether lipids synthesis take place in peroxisomes (Lodhi and Semenkovich 2014; Chornyi et al. 2023) and depletion of several e-PE species could suggest a defect of these organelles. The observed opposite changes in sphingolipid and e-PE contents align with the hypothesis of compensatory roles of these lipid classes in the regulation of the biophysical properties of cellular membranes (Jimenez-Rojo et al. 2020) While we observed both significant depletions of triglycerides and PEs, BLTP1 orthologs were typically linked to phospholipids in literature. Its yeast ortholog is interacting with an ethanolamine phosphotransferase(Toulmay et al. 2022), while the *C. elegans* LPD-3 is filled with phospholipids(Kang et al. 2025). Correspondingly, the cold shock sensitivity of *lpd-3* mutants could be rescued by phospholipid supplementation(Wang et al. 2022). In conclusion, our results suggest that non-vesicular lipid transport akin to ribosome biogenesis(Li et al. 2025) is essential for neocortical and cerebellar lamination.

## MATERIALS AND METHODS

### Protein model

A complete model for BLTP1 is not directly available from the AlphaFold repository as it provides overlapping fragments for proteins longer than 2,700 residues(Jumper et al. 2021; Varadi et al. 2022; Varadi et al. 2024). These were downloaded from the ftp site [https://ftp.ebi.ac.uk/pub/databases/alphafold] and a complete model was assembled with Swiss-PdbViewer(Johansson et al. 2012). Note that the swissprot entry sp|Q2LD37 used by AlphaFold with 5,005 residues is shorter than the MANE transcript (NM_001384125.1) that encodes a 5,091 residues protein. For BLTP1, 20 overlapping fragments are available: AF-Q2LD37-F1-model_v4.pdb.gz to AF-Q2LD37-F20-model_v4.pdb.gz. Fragments F1, F3, F5, F9, F15 and F20 were loaded each in their own layer and residues renumbered to match the actual residue number of the full-length sequence. Corresponding residues of various fragments were selected and used to superpose the fragments with the “fit selection tool”.

F3 onto F1: YTPAIKGQLLHVDATTSMQYRTLLEAEMLAFHINASY

(backbone rmsd 0.14 Angstroms)

F5 onto F3: REGHINLSGLQLRAHAMFSAEGLPLGSDSLEYAWLIDVQAGSLTAKV

(backbone rmsd 0.31 Angstroms)

F9 onto F5: IAAKLNIHRVHGQLRGLDTTDIGTCAITAIPFEKSKVLFTLEELDE

(backbone rmsd 1.17 Angstroms)

F15 onto F9: LTVVKCSIAKSQALYSAQRGLKTNNAAVFKVGAISINIP

(backbone rmsd 0.60 Angstroms)

F20 onto F15: PVDVVVYVRVQPSQIKFSCLPVSRVECMLKLPSLDLVFSSN

(backbone rmsd 1.26 Angstroms)

Then the following residues were selected in each layer: F1: 1-1110, F3: 1111-1734, F5: 1735-2159, F9: 2160-2863, F15: 2864-4080 and F20: 4081-5005 and a final composite model was assembled using the “create merged layer from selection (by layer)” tool. To assess the variants, we used the experimentally solved cryo-EM structure of the LPD-3 N-terminal region, the BLTP1 *C. elegans* ortholog (PDB entry 9CAP), generously shared prior publication by Sarah Clark(Kang et al. 2025). The level of structural conservations of residues not resolved in the cryo-EM structure were evaluated using the AlphaFold model of LPD-3 taken directly from the supplementary material of reference(Pandey et al. 2023). It was superposed on the human model selecting the pairs of structurally conserved variants (nematode N206 on human N216; P1246 on human P1631; P2354 on human P3050) using the fit backbone of selected residues option (12 atoms, rmsd (root mean square deviation 2.19 Angstroms). Of note, the sequence of the cryo-EM structure and AlphaFold model differ by two residues (for example E220 = E218, respectively). To avoid confusion, we kept the original residue numbering of the models and indicate which structure was used to identify equivalent residues (i.e cryo- or AlphaFold model). These models were used to evaluate the potential impact of missense variants (**Figure 1, Table S3**).

### Mouse husbandry

*Bltp1* knock-out first allele (tm1a) mice were ordered from the EMMA repository with the international strain name C57BL/6N-4932438A13Rik^tm1b(EUCOMM)Hmgu^/Ieg. In this strain, a targeting vector was inserted into *Bltp1* intron 9 and floxing exon 10 by homologous recombination in embryonic stem cells. Both FRT and LoxP sites allow for versatile remodeling (**Figure S1**)(Skarnes et al. 2011). All procedures were performed in accordance with protocols approved by the local relevant institutional authorities (canton Vaud veterinary office authorization VD3485; Direction Départementale de la Protection des Populations, article 17 of regulation number 1069/2009, reference number DDPP21 2022 01124). They complied with the European Convention for the Protection of Animals for Experimental and Scientific Purposes (ETS number 123) and the National Institutes of Health guide for the care and use of laboratory animals.

### Neuroanatomical characterization of adult mouse brains

To acquire brain samples derived from Emx1^Cre^*; Blpt1*^fl/fl^ and Emx1^Cre^*; Blpt1*^+/fl^, adult mice were first anesthetized, followed by brain dissection and fixation in 10% neutral buffered formalin. Neuroanatomical studies were carried out using two heterozygous *Bltp1* cKO male mice and the lone surviving homozygous *Bltp1* cKO male mouse. In this specific experiment, the heterozygous mice were used as controls, due to the lack of available littermate WT mice. Of note we do not observe differences between WT and heterozygous cKO embryos (see bwlow). Paraffin-embedded brain samples were cut at a 5μm thickness with a sliding microtome (Micom HM 450) to obtain brain sections at Lateral +0.60 mm, according to the Allen Mouse Brain Atlas(Collins et al. 2018). Sections were stained with 0.1% Luxol Fast Blue (Solvent Blue 38; Sigma-Aldrich) for myelin and 0.1% Cresyl violet acetate (Sigma-Aldrich) for neurons and scanned using the high-resolution digital slide scanner (Nanozoomer 2.0HT, C9600 series, Hamamatsu Photonics, Shizuoka, Japan) at 20× resolution.

A total of 40 brain morphological parameters, made of area and length measurements, were taken blind to the genotype using scripted routines and manual segmentation on ImageJ (https://imagej.net/software/fiji/). These 40 measurements encompassed 22 brain regions (**Table S4**): 1) the total brain area (TBA), the brain width (TB_Width), and heights (TB_Height1 at Bregma +0.86 mm and TB_Height2 at Bregma -1.34 mm); 2) the area of the temporal cortex (TCTX), the length of the secondary (M2) and primary (M1) motor cortices; 3) the height of the pons (pons); 4) the total cerebellar area (TCA), the internal granular layer of the cerebellum (IGL), the number of folia (Folia), and the area of the medial cerebellar nucleus (Med); 5) the area of the lateral ventricle (LV); 6) the area, the length and the height of the corpus callosum (cc); 7) the area of the thalamus (Th); 8) the area of the caudate putamen (CPu); 9) the area of the hippocampus (HP), the length of the radiatum layer of the hippocampus (Rad), the length of the oriens layer of the hippocampus (Or), the total internal length and area of the pyramidal cell layer of the hippocampus (TILpy), the length of the lacunosum moleculare layer (Mol), the area and the length of the granule cell layer of the dentate gyrus (DG); 10) the area of the fimbria of the hippocampus (fi); 11) the area of the anterior commissure anterior part (aca); 12) the area of the stria medullaris (sm); 13) the area of the fornix (f); 14) the area of the optic chiasm (opt); 15) the area of the ventromedial hypothalamus (VMH); 16) the area of the pontine nuclei (Pn); 17) the area of the substantia nigra (SN); 18) the area of the transverse fibers of the pons (fp); 19) the area and height of the cingulate cortex (Cg); 20) the area of the dorsal subiculum (DS); 21) the area of the inferior colliculus (InfC); and 22) the area of the superior colliculus (SupC).

### 3D and 2D neuroanatomical characterization of embryonic mouse brains

Pregnant female mice at 18.5 dpf were sacrificed and pups were dissected out of the uterus and placed on an ice-cold Petri dish. For male (N=12 WT, N=5 Emx1^Cre^*; Blpt1*^+/fl^, and N=4 Emx1^Cre^*; Blpt1*^fl/fl^) and female (N=4 WT, N=4 Emx1^Cre^*; Blpt1*^+/fl^, and N=5 Emx1^Cre^*; Blpt1*^fl/fl^) E18.5 embryos, the whole body was fixed in Bouin solution for 48 hours before undergoing either 2D or 3D histological processing, as described below.

Two embryonic heads per genotype group were removed and transferred to 70% ethanol then dehydrated in successive baths of ethanol of increasing concentration as previously described for HREM imaging(Montillot et al. 2023). After dehydration, samples were infiltrated at 4 °C with gentle rocking for seven days in JB-4 methacrylate plastic resin (Polysciences Europe GmbH, Germany) containing eosin B (Sigma-Aldrich). After the addition of a catalyst, the blocks were left to polymerize overnight at room temperature and baked at 95 °C for 48 hours, then cooled down overnight at 4 °C to ensure a hard texture prior sectioning. Serial face sectioning was done using 3 μm sections. By sequentially imaging the block face during the sectioning process, a comprehensive stack of accurately aligned thousands of images was acquired, producing the 3D structure of the sample. Three-dimensional morphometric analyses were performed using the SlicerMorph extension in 3D Slicer version 5.8(Rolfe et al. 2021) A total of 26 anatomical landmarks were manually placed on each embryonic brain model to capture key neuroanatomical features. To achieve dense surface correspondence across specimens, 601 pseudolandmarks were automatically generated using the Dense Correspondence Analysis (DeCAL) module (https://github.com/SlicerMorph/SlicerDenseCorrespondenceAnalysis)(Rolfe and Maga 2023). The resulting landmark configurations were subjected to Generalized Procrustes Analysis (GPA) to remove non-shape variation due to translation, rotation, and scale. Principal Component Analysis (PCA) was subsequently applied to the aligned coordinates to visualize and quantify major axes of shape variation among experimental groups.

The remaining embryonic brains were harvested, transferred to 70% ethanol, and manually embedded in paraffin using the following steps: three incubation baths in 70% ethanol for 30 minutes each, two baths in 95% ethanol for 30 minutes each, two baths in 100% ethanol for 45 minutes each, three baths in Histosol Plus for 1 hour each, and five baths in warm paraffin (60°C) for 30 minutes each, followed by incubation in warm paraffin overnight before casting in a mold. Brains were cut at a thickness of 5μm on a microtome (HM 450, Microm Microtech, France), such that we obtain sections matching planes at precise embryonic sagittal sections, as explained in (Nguyen et al. 2022). The sections were stained with 0.1% Cresyl violet acetate (Sigma-Aldrich) and scanned using Nanozommer 2.0HT, C9600 series at 20x resolution.

Each image was quality controlled to assess whether (i) the section is at the correct position, (ii) the section is symmetrical, (iii) the staining is of good quality, and (iv) the image is good quality. Only images that fulfilled all the quality control checks were processed. These quality control steps are essential for the detection of small to moderate neuroanatomical phenotypes, without which most neuro-anatomical phenotypes would be missed(Collins et al. 2019). Forty brain morphological parameters (area and length measurements) were taken for each sample at E18.5 (**Table S5 and S6** for female and male, respectively). Assessed embryonic brain regions included the total brain area, the cortices (motor, insular, somatosensory, retrosplenial granular and motor), the hippocampus, the genu of the corpus callosum, the internal capsule, the caudate putamen, the fimbria of the hippocampus, the anterior commissure, and the ventricles (lateral and third). Every aspect of the procedure was managed through a relational database using the FileMaker (FM) Pro database management system (detailed in ^20^). A list of histological parameters is also provided in **Figure 3**. Data were analyzed using two-tailed Student’s *t*-tests of equal variances.

### Immunofluorescence

Heads of E18.5 embryos were sectioned and put in 14ml ice-cold PBS. A piece of tail was kept for genotyping. The heads were then placed in tissue cassettes and incubated for 24h in PAXgene Tissue FIX reagent (PreAnalytiX, ref.765312), followed by an overnight incubation at 4°C in PAXgene Tissue Stabilizer reagent (PreAnalytiX, ref. 1075425). They were then rapidly washed 3x in 80% EtOH and put to dehydrate 2x 1hr in 80% EtOH, 2x 1hr in 95% EtOH, 1hr in 100% EtOH, 1hr30 in 100% EtOH, 1hr in 50% EtOH/50% Xylol, 1hr30 in 50% EtOH/50% Xylol, 1hr in Xylol, 1hr30 in Xylol and 2x 2hrs in paraffin. Heads were cut on the microtome to 4μm thickness onto microscope slides. Sections were dewaxed by three successive xylol baths and rehydrated by successive baths in 2x 100% EtOH, 95% EtOH, 80% EtOH, 70% EtOH and ddH2O. Antigen retrieval by heat-induced epitope retrieval (HIER) was performed on the slides by doing a quick wash followed by 10 minutes in the microwave with 0.01M citrate buffer pH6.0 and cooling at 4°C for 15 minutes to reach 60°C. Slides were then washed 3x 5 minutes in PBS and blocked 1hr at RT with PBS containing 2% normal goat serum and 0.3% Triton X-100 (T-PBS). Incubation with primary antibody were performed in T-PBS, first 2hrs at RT and then overnight at 4°C. Primary antibody references and dilutions are as follows: Rat anti-Ki67 SolA15 (Invitrogen 41-5698-80) 1/60 working dilution, Rabbit anti-Tbr1 (Abcam 31940) 1/200, Rabbit anti-PAX6 (Sigma AB2237) 1/400, Rat anti-EOMES (Tbr2) (Invitrogen 14-4875-82) 1/100, and Rabbit anti-cleaved Caspase-3 (Asp175) (Cell Signaling 9661) 1/100. Slides were then washed 3x 5 minutes in PBS and incubated in the dark at RT for 30 minutes in T-PBS with fluorescently labelled secondary antibodies Goat anti-rabbit Alexa488 (Invitrogen A11034) 1/500 or Goat anti-rat Alexa568 (Invitrogen A11077) 1/500, depending on host of primary antibody, as well as DAPI. Slides were washed 3x 5 minutes in PBS and mounted with Mowiol, kept in the dark 1-2hrs at RT and subsequently stored at 4°C. Slides were visualized under an upright fluorescence microscope.

### UK Biobank

The UK Biobank has enrolled about half a million British volunteers from the British population (Bycroft et al. 2018). Within the UK Biobank, we identified 1466 participants who are heterozygote for one of 326 *BLTP1* truncation variants. For those participants whose phenotypes were available, we compared the volumes of the hippocampus, the cortical-nucleus, the whole-brainstem (N carrier=189, N control=171) and the peripheral cortical grey matter normalized for head size (N carrier=183, N control=166).

### Lipidome profiling

Embryonic cortex (∼130 mg) was homogenized and frozen in RIPA lysis buffer containing protease inhibitors, using 110 µL per 10 mg of tissue. An aliquot of 25 µL of this homogenate was extracted with 125 µL of isopropanol(Medina et al. 2020) pre-spiked with an internal standard (IS) mixture containing 75 isotopically labeled lipid species (with multiple representatives per lipid class and varying fatty acid compositions). The resulting lipid extracts were analyzed by hydrophilic interaction chromatography coupled to electrospray ionization tandem mass spectrometry (HILIC ESI-MS/MS) in both positive and negative ionization mode, using a TSQ Altis LC-MS/MS system (Thermo Scientific), as previously described(Medina et al. 2023). The initial qualitative screen included more than 2,100 lipid species previously reported across diverse biofluids and tissue extracts. Following this screen, robustly detected species (across five replicates of pooled extract) were quantified in two separate runs using a dual-column setup. In total 429 lipid species belonging to five major classes of complex lipids (glycerolipids, glycerophospholipids, cholesterol esters, sphingolipids and free fatty acids) were quantified with high precision and specificity. Optimized lipid class-dependent parameters were used for data acquisition in timed selected reaction monitoring mode. Raw LC-MS/MS data were processed using the Trace Finder Software (version 4.1, Thermo Fisher Scientific). Lipid abundances are reported as the estimated concentrations based on single-point calibration using IS mixtures. Lipid Ontology Enrichment analysis was performed with the LION web application(Molenaar et al. 2019; Molenaar et al. 2023).

## COMPETING INTEREST STATEMENT

The authors do not have any competing interests to declare

## ACKNOWLEDGMENTS

We would like to thank Sarah Clark for sharing data prior to publication. This study made use of data generated by the International Mouse Phenotyping Consortium (IMPC). This work was supported by grants from the Swiss National Science Foundation (31003A_182632 and IZSTZ0_216615 to AR), the Lejeune Foundation (#1838-2019A to AR) and the Blackswan Foundation (to AR). BY is an INSERM investigator. Acquisition and use of the HREM equipment were supported by grants from the French National Research Agency (ANR-18-CE12-0009 to BY) and the European Union through the FEDER program (WDR to BY). We are grateful to the staff of the ImaFlow core facility (Biologie Santé Dijon BioSanD US58, 21079, Dijon, France), supported by Burgundy Regional Council (in particular Amandine Bataille and Audrey Geissler), and to the CCuB for technical support and management of the computing platform. We are grateful to the students of the NeuroGeMM Laboratory (in particular Emeline Richter, Elisa Mischler, Lucile Tonneau, Sylvie Nguyen, and Charlène Parisis) for their involvement in the mouse neuroanatomical studies. Some authors are members of the European Reference Network on Rare Congenital Malformations and Rare Intellectual Disability ERN-ITHACA, which is funded by the European Union under agreement N°101156387. UK Biobank has approval from the NorthWest Multicentre Research Ethics Committee as a Research Tissue Bank. All participants signed a broad informed consent form. Data were accessed under application 16389. The funders had no role in study design, data collection and analysis, decision to publish, or preparation of the manuscript.

## AUTHORS’ CONTRIBUTIONS

LG, FA, FTMT, DCK, MTP, KVT, LZR, LW, KS, MAD, CO, JR, MM, ACJ, JH, MK, APC, BP, EW, OK, AB, JH, JHC, SA, MFS and SB collected clinical information and DNAs, sequenced and analyzed exomes and/or genomes. NG 3D-modeled missense variants. JC, SCC, CR, SBA, CA, OM, YE and BY engineered and phenotyped animal models. PR, CA and ZK analyzed UK Biobank data. HGA, NG, FV, MA and JI performed the lipidomics experiments and analyzed its results. AR conceived the study and wrote the manuscript. All other authors read and approved the manuscript.

## FIGURE LEGENDS

**Figure S1.**
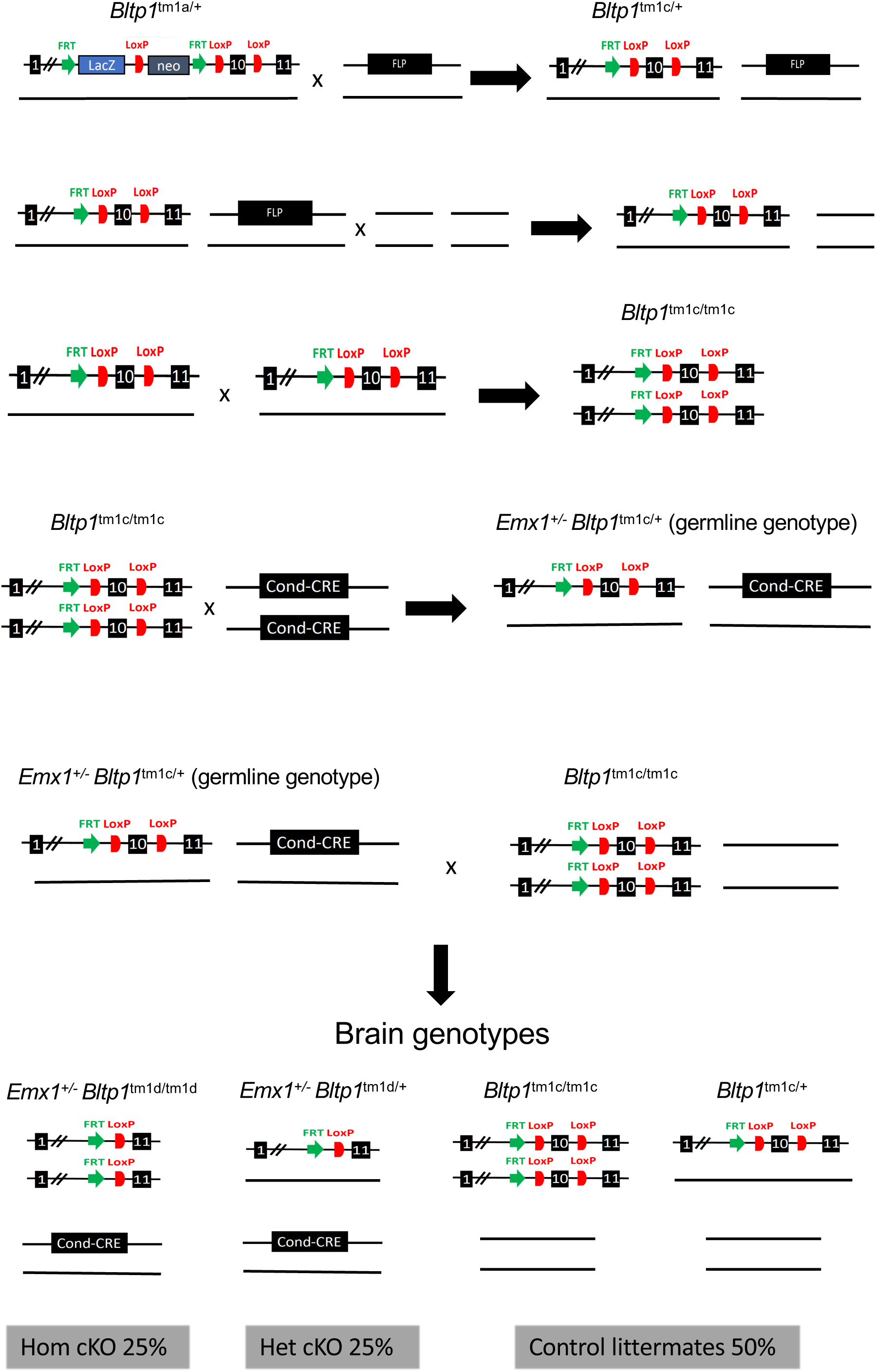
Breeding scheme. Schematic presentation of the consecutive breeding crosses implemented to produce cKO (*Emx1-Cre; Bltp1^fl/fl^*) individuals (adapted from reference (Skarnes et al. 2011)).

**Figure S2.**
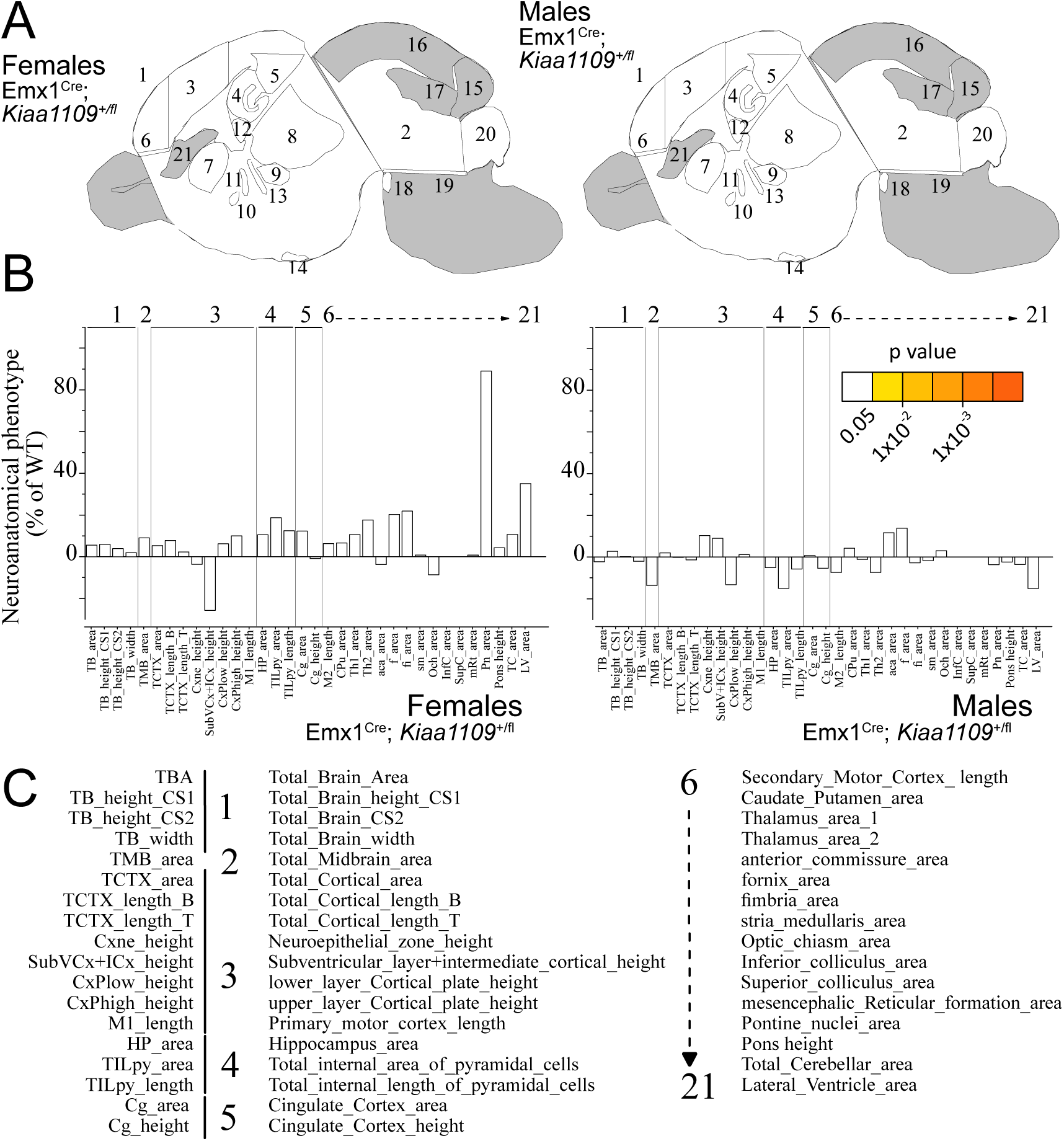
***Bltp1* is haplosufficient in corticogenesis.** (**A**) Schematic illustration of a mouse sagittal brain section at embryonic day 18.5 (E18.5). Numbers identify the different brain regions assessed for area and length differences between cKO (*Emx1-Cre; Bltp1^+/fl^*) and control littermates in females (**left**) and males (**right**). White indicates a p-value > 0.05, grey shows not enough data to calculate a p-value. (**B**) Histograms depicting the size changes observed in female (**left**) and male (**right**) cKO embryos relative to littermate control embryos (set as 0) for each of the measured parameters (listed in **Tables S5-S6**). (**C**) List of parameters measured with full name. Statistical analyses were performed with R, using two-tailed Student’s *t*-tests of equal variances.

**Figure S3.**
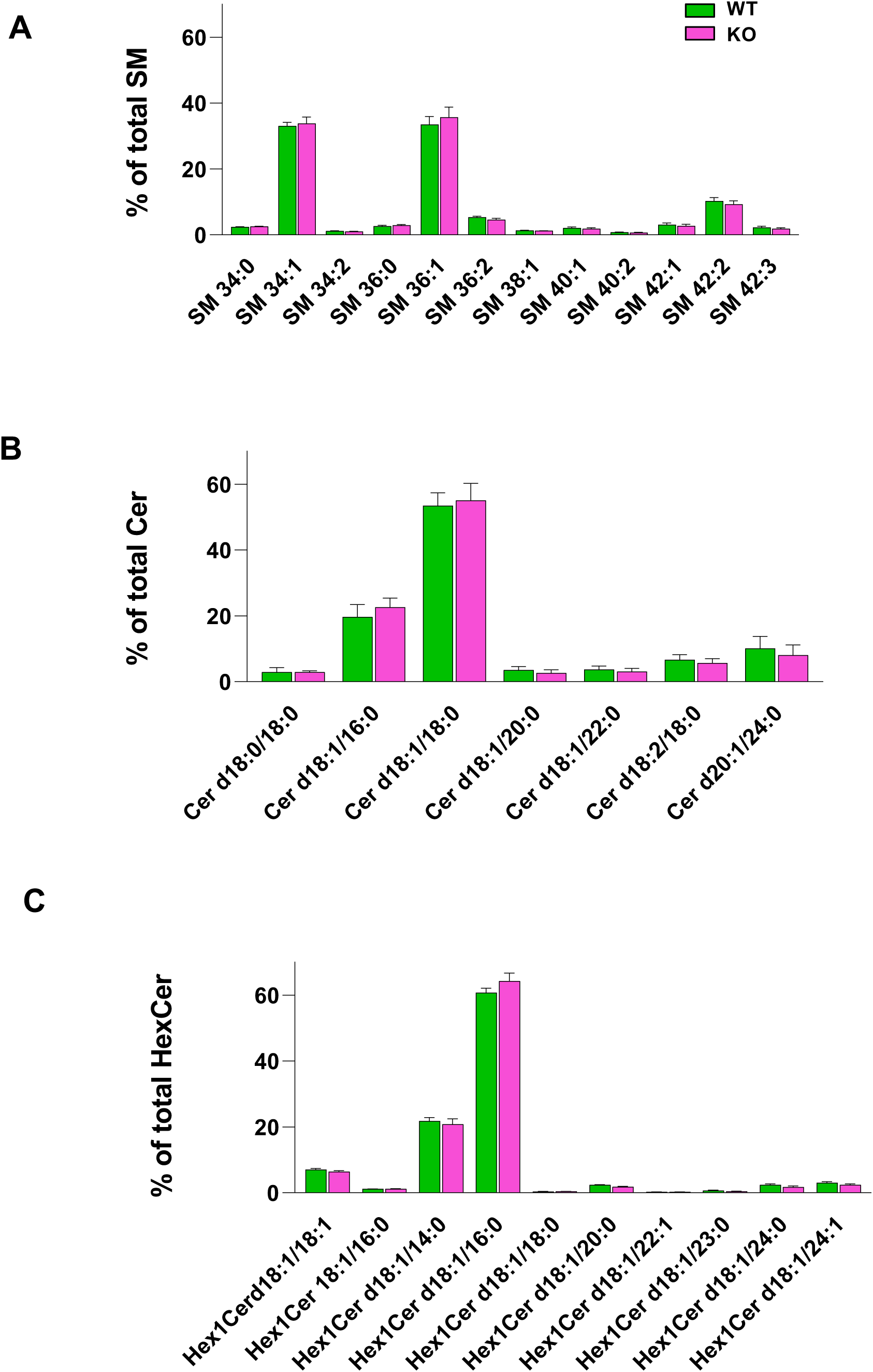
Sphingolipid species distribution. Relative abundance of individual SM **(A)**, Cer **(B)** and HexCer **(C)** species in WT (green) and cKO (pink) E18.5 cortexes expressed as a percentage of the respective class.

**Figure S4.**
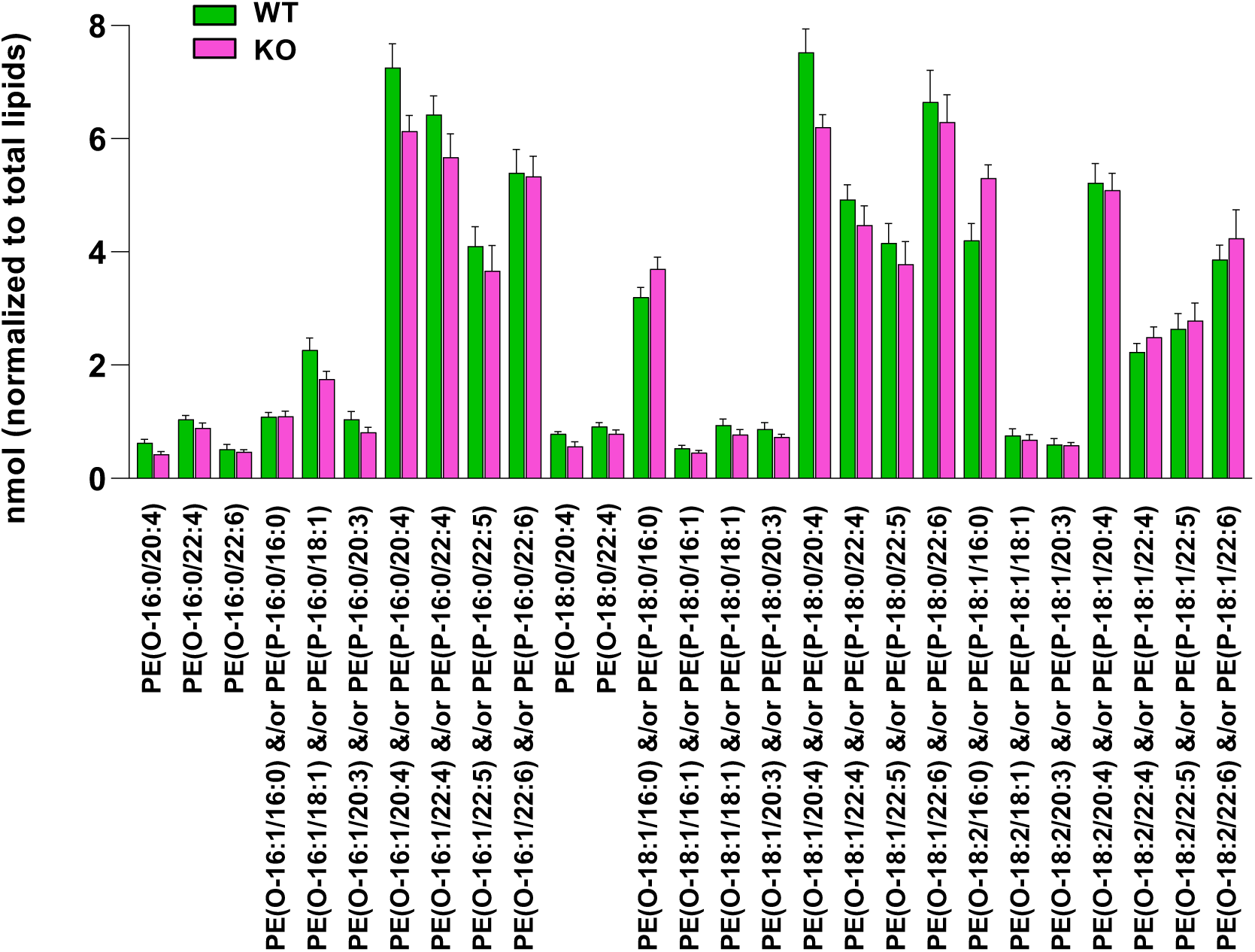
Alterations of individual e-PE species. Molar abundance of individual e-PE species in WT (green) and cKO (pink) E18.5 cortexes.

**Table S1: genotypes and phenotypes of published Alkuraya-Kučinskas syndrome individuals**

**Table S2: genotypes and phenotypes of novel Alkuraya-Kučinskas syndrome individuals**

**Table S3: evaluation of Alkuraya-Kučinskas syndrome-associated missense variants**

**Table S4: measurements of 40 parameters encompassing 22 brain regions in adult mice**

**Table S5: measurements of 40 parameters encompassing 17 brain regions in E18.5 females**

**Table S6: measurements of 40 parameters encompassing 17 brain regions in E18.5 males**

**Table S7: measurements of 394 lipid metabolites in cortex**

